# Distinct Solubility and Cytotoxicity Regimes of Paclitaxel-Loaded Cationic Liposomes at Low and High Drug Content Revealed by Kinetic Phase Behavior and Cancer Cell Viability Studies

**DOI:** 10.1101/154948

**Authors:** Victoria M. Steffes, Meena M. Murali, Yoonsang Park, Bretton J. Fletcher, Kai K. Ewert, Cyrus R. Safinya

**Author notes:** Present address: Department of Materials Science and Engineering, Pennsylvania State University, University Park, PA 16802, USA.

## Abstract

Lipid-based particles are used worldwide in clinical trials as carriers of hydrophobic paclitaxel (PTXL) for cancer chemotherapy, albeit with little improvement over the standard-of-care. Improving efficacy requires an understanding of intramembrane interactions between PTXL and lipids to enhance PTXL solubilization and suppress PTXL phase separation into crystals. We studied the solubility of PTXL in cationic liposomes (CLs) composed of positively charged 2,3-dioleyloxypropyltrimethylammonium chloride (DOTAP) and neutral 1,2-dioleoyl-*sn*-glycero-3-phosphatidylcholine (DOPC) as a function of PTXL membrane content and its relation to efficacy. Time-dependent kinetic phase diagrams were generated from observations of PTXL crystal formation by differential-interference-contrast microscopy. Furthermore, a new Synchrotron small-angle x-ray scattering in situ methodology applied to DOTAP/DOPC/PTXL membranes condensed with DNA enabled us to detect the time-dependent depletion of PTXL from membranes by measurements of variations in the membrane interlayer and DNA interaxial spacings. Our results revealed three regimes with distinct time scales for PTXL membrane solubility: hours for > 3 mol% PTXL (low), days for ≈ 3 mol% PTXL (moderate), and ≥ 20 days for < 3 mol% PTXL (long-term). Cell viability experiments on human cancer cell lines using CL_PTXL_ nanoparticles (NPs) in the distinct CL_PTXL_ solubility regimes reveal an unexpected nonmonotonic dependence of efficacy on PTXL content in NPs delivered at short time scales (hours) after liposome hydration, where we see two distinct high-efficacy regimes at low (< 2 mol%) and high (9 mol%) drug loading. These newly identified high-efficacy regimes flank the membrane solubility limit (≈ 3 mol%, where efficacy declines), which has been the focus of most previous physicochemical studies (and clinical trials) of PTXL-loaded CLs. At longer times scales (days), CL_PTXL_ NPs with ≥ 3 mol% PTXL lose efficacy while formulations with 1–2 mol% PTXL maintain high efficacy. Our findings underscore the importance of understanding the relationship of the kinetic phase behavior and physicochemical properties of CL_PTXL_ NPs to efficacy.

## Introduction

The landmark discovery that paclitaxel (PTXL), derived from the Pacific Yew tree, suppresses cell division in tumors [1], has resulted in the ongoing worldwide effort to develop efficient synthetic carriers of PTXL for cancer chemotherapy [2]. PTXL is a hydrophobic molecule (Figure 1a,b) known to inhibit mitosis by stabilizing microtubules (upon binding a specific hydrophobic pocket on the β-tubulin subunit), thereby obstructing chromosome capture and segregation during metaphase and subsequently activating apoptotic signaling pathways that lead to cell death [3–8]. PTXL is among the most common drugs used to treat ovarian, breast, lung, pancreatic, and other cancers and is included in the World Health Organization’s List of Essential Medicines [9–13]. PTXL is commonly administered in the form of Taxol^®^, where it is solubilized for delivery in Kollifor-EL (formerly Cremophor EL), which causes hypersensitivity reactions and delivers PTXL non-discriminately throughout the body [14–16]. In 2005, nanoparticle albumin-bound PTXL was approved by the FDA (Abraxane^®^); this formulation is considered to have fewer adverse reactions than Taxol, although there are mixed reports on whether it improves survival outcomes [17–19].

**Figure 1.**
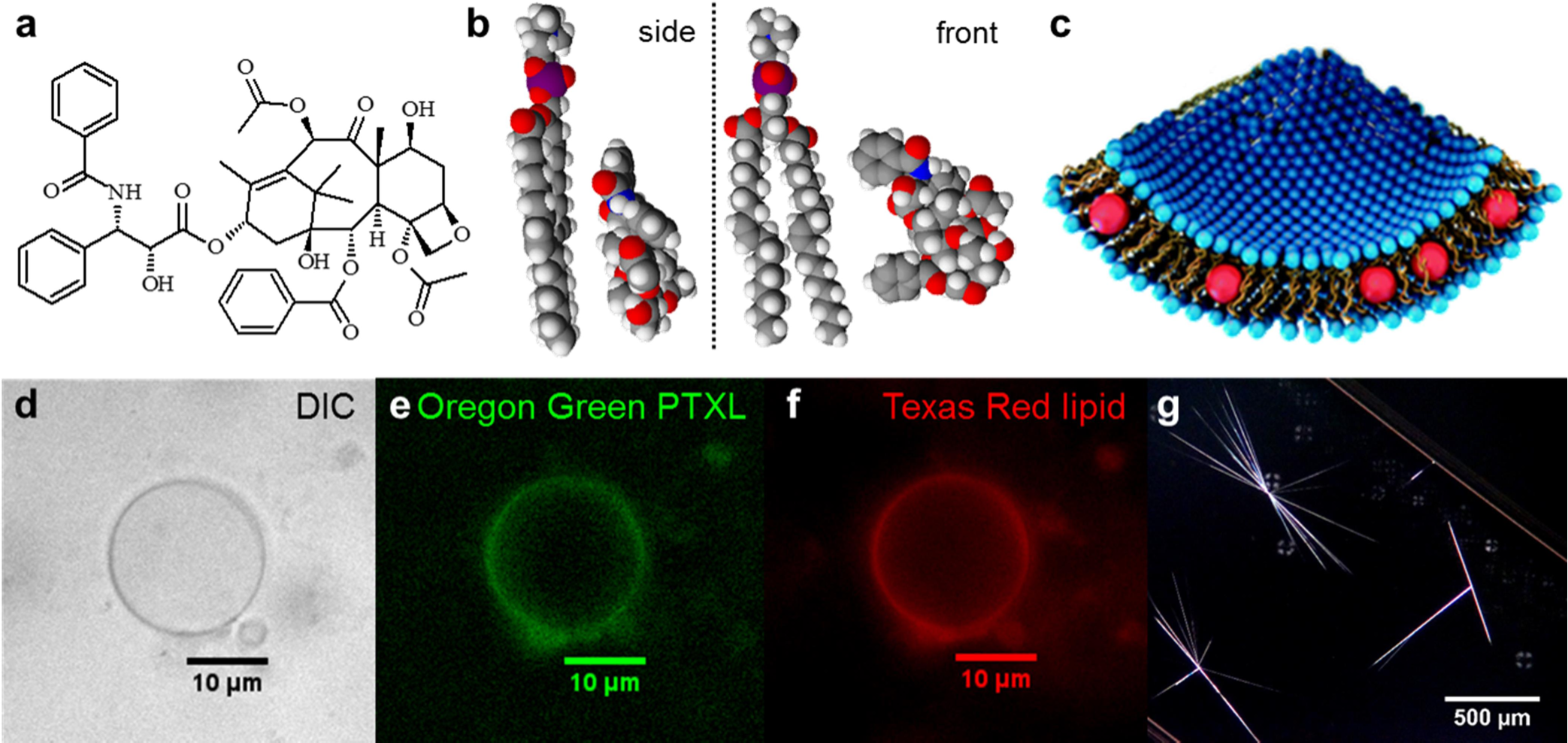
PTXL-containing liposomes from the molecular to micrometer scale. (a–c) a molecular look at the PTXL–lipid system: a) the chemical structure of PTXL, b) space filling molecular models for DOPC and PTXL viewed from the side and the front, c) schematic drawing of a liposome with hydrophobic molecules (red spheres, representative of PTXL) embedded within the membrane. (d–f) Microscopy images of a singular PTXL-containing liposome (composed of a 90:10:5:7.1 mole ratio of DOTAP:DOPC:OregonGreen-PTXL:TexasRed-DHPE), demonstrating colocalization of PTXL with the lipid bilayer: d) differential-interference-contrast image, e) green fluorescence due to Oregon Green-conjugated PTXL, f) red fluorescence from the Texas Red–DHPE lipid label. (g) Polarized optical microscopy image of PTXL crystals that have phase separated from liposomes (5:92:3 initial mole ratio of DOTAP/DOPC/PTXL).

To harness the intrinsic therapeutic capacity of PTXL, it needs to be delivered by a nontoxic carrier. Liposomal nanoparticles (NPs) are versatile drug delivery agents due to their ability to sequester multiple distinct therapeutic molecules, including long and short nucleic acids (electrostatically condensed with membranes) [20], as well as hydrophilic and hydrophobic drugs in different forms (solubilized, crystallized, surface-conjugated) [21]. Because each therapeutic drug molecule possesses unique physical and chemical characteristics, its liposomal carrier must be tailored to these properties to achieve efficient delivery [22].

Studies suggest that longer retention times of an anticancer drug within circulating liposomes leads to greater accumulation of the drug at tumor sites. Much of this work has been done using doxorubicin and other drugs that can be loaded efficiently in the interior aqueous pocket of liposomes (e.g. Doxil and Myocet) [23]. To prolong retention of these drugs in liposomes, the current approach is to increase the rigidity of the membrane and thereby reduce its permeability to hydrophilic drugs, by employing lipids with high melting points (e.g. with saturated tails) and including cholesterol [24,25].

Hydrophobic drugs, on the other hand, are solubilized by and reside directly in the nonpolar (hydrocarbon chain) region of the membrane (Fig. 1b,c). These drugs, which include PTXL, are quickly expelled from membranes consisting of saturated lipid tails or those that have a high concentration of cholesterol [26–31]. This observation indicates that the maximum loading and residence time of hydrophobic drugs within liposomes are particularly sensitive to the liposome composition. A key concern is that hydrophobic drugs will leach out of liposomes quickly in vivo because they reside at the particle boundary rather than the interior, and will subsequently bind to plasma proteins with hydrophobic pockets acting as ‘lipid sinks’ [21,25,32].

Various studies indicate that liposome–PTXL formulations exhibit lower toxicity compared to Taxol^®^, may increase the maximum tolerated drug dose, and may improve biodistribution [28,33–36]. One liposomal formulation of PTXL is approved in China (Lipusu^®^) [37,38], while others are in clinical trials. Composition information for the Lipusu formulation is not publically available. LEP-ETU is in Phase II trials in the United States; it is an anionic lipid-based carrier composed of the neutral lipid DOPC (1,2-dioleoyl-sn-glycero-3-phosphatidylcholine), cholesterol, and cardiolipin (90:5:5 mole ratio) with an additional 3 mol% PTXL [39]. EndoTAG-1 is in Phase III trials in Taiwan and has a cationic liposome structure consisting of the univalent cationic lipid 2,3-dioleyloxypropyltrimethylammonium chloride (DOTAP), DOPC, and PTXL (50:47:3 mole ratio) [26]. Other types of PTXL-containing liposomes (e.g. PEGylated, antibody-targeted) have shown some promising early results, but have not progressed to clinical trials [28,33,40–43].

Straubinger has reported that 3 mol% is the solubility limit for PTXL in liposomes, above which PTXL rapidly precipitates [44]. However, there are few studies that systematically evaluate this drug loading parameter as a function of lipid and membrane properties and none that demonstrate that liposomes at this limit are the most efficient at actual drug delivery. Studies from various groups have investigated the behavior of PTXL in different types of membranes—cationic, anionic, neutral, cholesterol-containing, saturated, and PEGylated—but these studies are few in number and together form a very fragmented and sometimes contradictory picture [27–30,42,45]. What is lacking is a comprehensive study of how the physicochemical properties of liposome–PTXL particles, including their time-dependent phase behavior, correlate to PTXL membrane content for optimal drug delivery and efficacy (i.e. ability to induce cancer cell death).

Cationic liposomes (CLs) are of particular interest because the electrostatic attraction between positive particles and the negatively charged sulfated proteoglycans (components of the glycocalyx which covers cells) leads to cell binding. Studies suggest that positively charged particles may achieve some cancer-specific targeting because the tumor neovasculature is more negatively charged than other tissues in the body and will therefore accumulate cationic particles at a higher concentration [40,46–48]. This assertion was the basis for choosing the lipid mixture of EndoTAG-1 as a starting point in our investigation of PTXL–lipid interactions.

In the work reported here, we generated kinetic phase diagrams characterizing the time-dependent phase separation and crystal formation of PTXL as a function of PTXL content in DOTAP/DOPC membranes. The phase diagrams were generated from direct observations of PTXL crystallization using differentialinterference-contrast (DIC) microscopy. To corroborate this data, we used high-resolution synchrotron small-angle x-ray scattering (SAXS), which unambiguously identified the PTXL crystals by their characteristic diffraction peaks. Furthermore, SAXS allowed us to perform in situ measurements of variations in the interlayer spacing as a function of time in multilamellar, onion-like complexes of cationic DOTAP/DOPC/PTXL membranes condensed with DNA, confirming the depletion of PTXL from the membranes upon crystallization.

The kinetic phase diagrams show a solubility threshold at 3 mol% PTXL content: below 3 mol%, PTXL exhibits long-term solubility (≥ 20 days) in unsonicated CLs, whereas above 3 mol%, the drug crystallizes within the first day following hydration. PTXL remained soluble in CLs on a time scale of days when incorporated at 3 mol%. The duration of PTXL solubility in CL_PTXL_ NPs consisting of small (< 200 nm diameter) unilamellar liposomes (produced by sonication) is shorter than in unsonicated uni- and multi-lamellar liposomes with a broad distribution of larger sizes (average diameter ≈ 800 nm) over the whole range of PTXL contents tested.

While previous studies seem to empirically choose one or two PTXL–liposome formulations based on physical characterization alone to push forward directly into animal testing, our study breaks from that approach and instead attempts to correlate the extent of biological response to the liposome properties. Thus, we assessed efficacy *in vitro* for a series of CL_PTXL_ NPs with varying PTXL content in the CL membranes (at fixed total PTXL concentration in solution) by measuring human cancer cell survival. We observed new drug delivery behavior dependent both on the PTXL content (i.e., PTXL loading) and the timing of drug delivery after liposome hydration (at short (hours) vs. long (days) times). CL_PTXL_ NPs that exhibit long-term solubility (PTXL content ≤ 2 mol%) show the highest efficacy (extent of cell death), whether delivered hours or days after liposome hydration. Surprisingly, CL_PTXL_ NPs containing 9 mol% PTXL exhibited greater efficacy than NPs with 3–7 mol% PTXL when the particles were applied to cells within a few hours of hydration. This effect disappears over time, as PTXL phase separates and crystalizes. Thus, whereas previous studies (including clinical trials) of CL-based carriers have exclusively focused on 3 mol% PTXL content (i.e. near the CL solubility limit) [33,39,49], our study has revealed two distinctly new PTXL composition regimes, below and above 3 mol% PTXL content, where CL_PTXL_ NPs exhibit improved efficacy. Furthermore, our results demonstrate that the time of NP delivery after liposome hydration is a critical parameter affecting efficacy.

## Materials and Methods

### Materials

Lipid stock solutions of DOPC and DOTAP in chloroform were purchased from Avanti Polar Lipids. PTXL was purchased from Acros Organics and dissolved in chloroform at 10.0 mM concentration. CellTiter 96^®^ AQueous-One Solution Cell Proliferation Assay was obtained from Promega. Paclitaxel– Oregon Green^®^ 488 Conjugate, Texas Red–DHPE, and glycerol monooleate (GMO) were purchased from Thermo Fisher Scientific as powders and dissolved in chloroform to 190 μM, 81 μM and 10 mM concentrations, respectively.

### Cell Culture

The human cell lines PC3 (ATCC number: CRL-1435; prostate cancer) and M21 (melanoma) were gifts from the Ruoslahti Lab (Burnham Institute, La Jolla). M21 cells are a subclone that was derived in the laboratory of Dr. Ralph Reisfeld (Scripps Institute, La Jolla) from the human melanoma line UCLA-SOM21, which was originally provided by Dr. D. L. Morton (UCLA, Los Angeles). Cells were cultured in DMEM (Invitrogen) supplemented with 10% fetal bovine serum (Gibco) and 1% penicillin/streptomycin (Invitrogen). Cells were passaged every 72 h to maintain subconfluency and kept in an incubator at 37°C in a humidified atmosphere containing 5% CO_2_.

### Liposome preparation

Mixed solutions of lipid and PTXL were prepared in chloroform:methanol (3:1, v/v) in small glass vials at a total molar concentration (lipid + PTXL) of 1 mM for cell experiments, 5 mM for DIC microscopy, and 20 mM for x-ray experiments. Individual stock solutions were combined according to the desired molar composition; typically, the cationic lipid DOTAP content remained constant for comparison, while neutral DOPC was exchanged for PTXL as the amount of PTXL was varied. The organic solvent was evaporated by a stream of nitrogen for 10 min and dried further in a vacuum (rotary vane pump) for 16 h. The resulting lipid/PTXL films were hydrated with high-resistivity water (18.2 MΩ cm) to the aforementioned concentrations. Immediately thereafter, suspensions were agitated with a tip sonicator (Sonics and Materials Inc. Vibra Cell, set to 30 Watt output) for 7 min to form small unilamellar vesicles (“sonicated liposomes”).

### Dynamic Light Scattering

A Malvern Zetasizer Nano ZS was used to measure the average size of the liposomes. A total of 100 μL of a 5 mM stock suspension was diluted with 900 μL of DMEM to mimic the salt conditions of the liposomes in solution when they are added to cells. This solution was loaded into a DLS cuvette (Malvern DTS1070) and measured. The liposome diameter is reported as the average ± standard deviation of 3 measurements.

### Polarized optical microscopy

Liposome samples at a concentration of 5 mM were loaded into flat microslides (VitroCom) via capillary action and sealed on both ends with 5-min epoxy glue. Samples were observed under crossed polarizers on a Nikon Optiphot microscope with a 5× objective.

### DIC microscopy

Samples prepared at 5 mM concentration were mixed manually after hydration by agitating the vial to ensure homogenous mixing. The sample solutions were stored at 37°C for the duration of the experiment. At predetermined times, 2 μL aliquots were withdrawn, placed on microscope slides, covered by a coverslip kept in place by vacuum grease, and imaged at 10 or 20× magnification on an inverted Diaphot 300 (Nikon) microscope. The samples were first imaged within minutes of adding water to the dried lipid films, then every 2 h until 12 h, every 12 h until 72 h, and daily thereafter until PTXL crystals were observed or the entire sample was used up.

### Fluorescence microscopy

Liposomes were prepared as described above at 5 mM total concentration, except for the incorporation of two fluorophores, one for lipid and one for PTXL. The liposome composition was DOTAP:DOPC:OregonGreen–PTXL:TexasRed–DHPE=90:10:5:7.1 (molar ratio). An aliquot of 2 μL of this solution was placed between glass coverslip and slide and sealed in with vacuum grease. The solution was imaged with a Nikon Diaphot 300 equipped with a Nikon 1.4 NA 60× Plan Apo DIC Objective and a PCO Sensicam QE CCD camera.

### X-ray scattering

X-ray scattering experiments were performed at the Stanford Synchrotron Radiation Lightsource (Beamline 4-2) with an x-ray energy of 9 keV and a sample–detector distance of 1.2 m. Lipid/PTXL films were prepared as described for liposome preparation (30 mol% DOTAP, 70–x_PTXL_ mol% DOPC). After hydration (to 20 mM, i.e., 6 mM DOTAP), lipid suspensions were placed in a bath sonicator for 30 min and then stored at room temperature. To prepare samples for x-ray scattering, aliquots of the liposome suspensions were mixed in 1.5 mm quartz capillaries (Hilgenberg) with calf thymus DNA solution (5 mg/mL in water) at a lipid/DNA charge ratio of 1.5. Samples were centrifuged in a capillary rotor in a Universal 320R centrifuge (Hettich, Germany) at 14,000 *g* for 30 min and the condensed lipid–DNA phase pelleted to the bottom of the x-ray capillary along with any PTXL crystals already present. Condensation of liposomes into CL–DNA complexes is expected to introduce confinement of crystal growth, which would complicate crystal detection by scattering. Thus, CL–DNA complexes were prepared immediately prior to measurement for each time point for each liposome composition under investigation, with the expectation that complex formation and centrifugation would break up newly formed crystals and disperse them heterogeneously into the CL–DNA pellet.

Concentrated lipid solutions with high PTXL content *without DNA* also produced opaque white pellets when centrifuged. X-ray scattering confirmed that these contained phase-separated PTXL. These results, shown in Figure 4a, are for samples using GMO as the neutral lipid in place of DOPC. The lipid only sample was DOTAP:GMO (50:50 mole ratio), while the PTXL sample had the composition DOTAP:GMO:PTXL (50:44:6 mole ratio).

### Cell viability assays

Cells were plated in 96-well plates at a density of 5,000 cells/well. Cells were incubated overnight to adhere to the plate. Liposome suspensions were diluted in DMEM to reach the desired concentration of PTXL. The cell culture medium was then manually removed from the wells with a pipette (rather than a vacuum aspirator; to ensure that cells were not removed unintentionally) and replaced with 100 μL of the liposome suspension. After incubation for 24 h, the liposome-containing medium was removed manually with a pipette and replaced with supplemented DMEM. After incubation for another 48 h, the cell viability was measured with the CellTiter 96^®^ AQueous-One Solution Cell Proliferation Assay (Promega). The assay solution was diluted 6-fold with DMEM and 120 μL of this solution were added to each well. The absorbance at 490 nm was measured with a plate reader (Tecan M220) after 1 h of incubation as per the assay instructions. Each data point is the average of four identically treated wells and reported as a percentage of the viability of untreated cells. The incubation times in this procedure are based on previous experiments by L. Wilson (UCSB) that were reproduced in our lab and showed that the viability of cells treated with PTXL relative to control cells decreases over time until reaching a plateau at around 72 h [4].

In viability experiments comparing CL_PTXL_ NPs of varying PTXL content, the total concentration of PTXL was kept constant (at the approximate IC-50, see text). This results in varying lipid concentrations. For example, delivering a fixed PTXL concentration of 40 nM in a volume of 100 μL at 1 and 9 mol% PTXL results in a total lipid concentration of 4 and 0.44 μM, respectively.

To assess the effect of time after hydration on the efficacy of CL_PTXL_ NPs, seven vials with CL/PTXL films were prepared as described above for each of the formulations (50 mol% DOTAP, 50–x_PTXL_ mol% DOPC) under investigation. These films were hydrated and sonicated over the course of 10 d (0, 1, 2, 4, 6, 8, and 10 d before addition to cells). On the day the CL_PTXL_ NPs were added to cells, each suspension was first diluted in DMEM to a final PTXL concentration of 22.5 nM for PC-3 cells and 65.0 nM for M21 cells. The solutions were then used in the cell viability assay as described above.

### Results and Discussion

#### Microscopy Characterization of PTXL-Loaded Cationic Liposomes

As described in the Introduction, hydrophobic drugs are expected to partition into the membrane of liposomes. To visualize this partitioning for PTXL in CL membranes, we carried out fluorescence colocalization experiments. Figure 1 (d-f) shows representative images of the results: part (d) displays a DIC optical image of a giant cationic liposome (composed of DOTAP:DOPC:OregonGreen–PTXL:TexasRed–DHPE at a 90:10:5:7.1 mole ratio); part (e) shows the green fluorescence from the Oregon Green^®^ 488–PTXL conjugate; part (f) shows the red fluorescence due to lipid label TexasRed–DHPE. This series of images demonstrates that fluorescent PTXL and lipid are colocalized and concentrated along the outer edge of the liposome, consistent with the PTXL conjugate residing within the hydrophobic region of the bilayer.

For the *quantitative* study of the stability of PTXL in CL formulations, the Oregon Green^®^ 488–PTXL conjugate is not a suitable PTXL analog because its chemical properties are significantly altered by the addition of the fluorophore. For example, while the conjugate can be used to image microtubules, it is applied to cells at a much higher concentration than lethal doses of unaltered PTXL, suggesting it is much less toxic. In addition, the fluorophore-conjugated PTXL is more water soluble than PTXL and does not readily phase-separate from the CL membranes to form crystals. We therefore used the unaltered form of PTXL (Figure 1a,b) for the remainder of our studies.

Various groups have reported 3 mol% PTXL as the solubility limit in liposomes [39,44]. Beyond this limit, PTXL phase-separates and forms characteristic needle-shaped crystals. Figure 1g shows a low-magnification polarized optical micrograph displaying such crystals in a CL_PTXL_ sample days after preparation. While previous studies have largely relied on multi-step HPLC experiments to monitor drug loading and retention over time [28,29,41], we set out to establish a DIC microscopy-based experimental protocol that would allow us to determine the PTXL stability in novel lipid formulations with higher throughput.

Crystal nucleation and growth theory and experimental evidence suggest that the rate-limiting step in PTXL crystallization is nucleation, with crystal growth occurring on a much faster time scale thereafter. Thus, direct observation of PTXL crystals using DIC microscopy is an appropriate technique to assess CL_PTXL_ stability on the relevant time scale of hours to days. Figure 2 provides examples of DIC micrographs of CL_PTXL_ liposomes before and after PTXL crystallization. The pictured samples consisted of 30 mol% DOTAP, 1.25–5 mol% PTXL, and the remainder DOPC. Two methods of sample preparation were used: unsonicated liposomes as formed spontaneously upon hydration, which are larger and display a broad size distribution, and sonicated liposomes, which display a more monodisperse distribution of smaller sizes.

**Figure 2.**
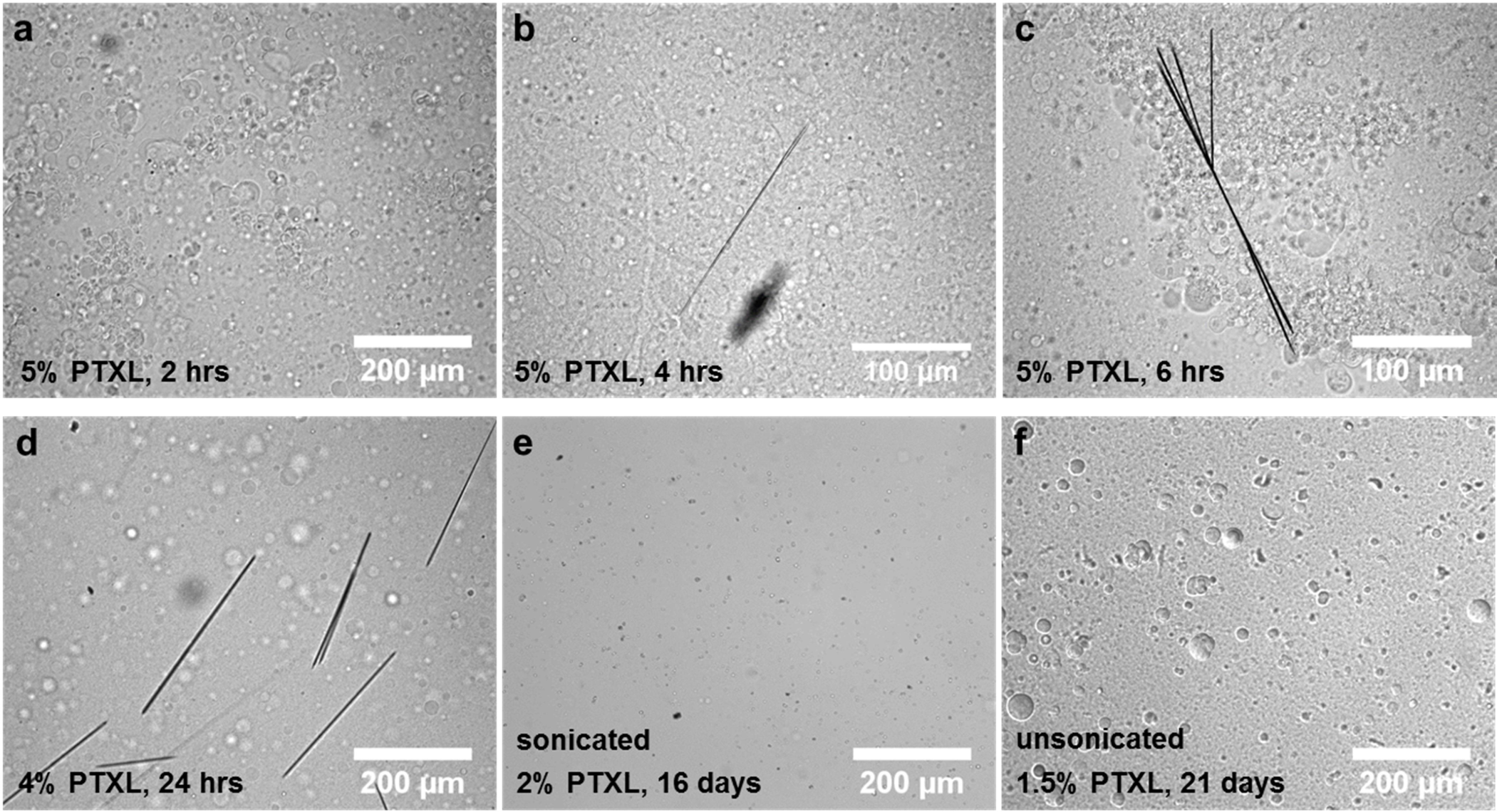
DIC microscopy images of liposomes and crystallized PTXL. Unless otherwise specified, samples are unsonicated liposomes made of DOTAP, DOPC and PTXL (30:70–x_PTXL_:x_PTXL_ mole ratio). (a–c) Images of liposomes with x_PTXL_=5 taken a) 2 h, b) 4 h, and c) 6 h after hydration; crystal formation is evident at 4 h, indicating the short period of stability at high PTXL contents. (d) Image of a sample with x_PTXL_=4, showing PTXL crystals at 24 h after hydration. (e and f) Images of samples that exhibit longer-term PTXL solubility: e) x_PTXL_=2, sonicated sample with an average particle size < 200 nm, and f) x_PTXL_=1.5, unsonicated sample composed of larger multilamellar vesicles (≈ 800 nm average diameter).

Figure 2 (a-c) displays images obtained for a sample in which the PTXL crystallizes in a matter of hours. The image shown in part (a) was taken 2 h after liposome hydration and shows no evidence of phase separation and crystal formation, while the images in parts (b) and (c) show PTXL crystals and an axialite bundle of crystals at 4 and 6 h, respectively. At lower PTXL content (4 mol%), CL_PTXL_ NPs are stable for longer periods, with PTXL crystals appearing between 12 h and 24 h (Figure 2d). The images in Figure 2e (2 mol% PTXL) and 2f (1.5 mol% PTXL) contrast the two sample preparation methods (sonicated (2e) and unsonicated (2f)) for two formulations at even lower PTXL content. These CL_PTXL_ NPs exhibited long term stability, with no crystals detectable by DIC microscopy even 16 days (Figure 2e) and 21 days (Figure 2f) after hydration. As is evident in the DIC images, unsonicated samples contain a broad distribution of particle sizes, up to tens of micrometers. This was confirmed by dynamic light scattering (DLS) measurements on CL_PTXL_ NPs at 30 mol% DOTAP but containing only 1 mol% PTXL: the average diameter of NPs for the sonicated sample was 180±20 nm, whereas the NPs in the unsonicated sample had an average diameter of 810±70 nm. Only a few liposomes at the upper limit of the size distribution of sonicated CL_PTXL_ NPs can be resolved by DIC microscopy (Figure 2e).

#### Kinetic Phase Behavior of PTXL-Loaded Cationic Liposomes

The time-dependent phase diagrams of unsonicated and sonicated PTXL-loaded CLs (DOTAP:DOPC:PTXL, 30:70-x:x mole ratio), as mapped out by DIC microscopy, are shown in Figure 3a and 3b, respectively. Blue color indicates that PTXL remained solubilized in membranes, i.e., no crystals were observed. Pink color indicates the time point at which PTXL crystals were observed. (Time points after the first observation of crystals were marked with pink color even if no further samples were assessed, since crystallization from the membrane is irreversible.) For example, in the unsonicated sample containing 5 mol% PTXL (DOTAP:DOPC:PTXL 30:65:5 mole ratio), crystals were first observed at the 4 h time point, meaning that PTXL crystallized between 2 and 4 h after hydration (Figure 3a).

**Figure 3.**
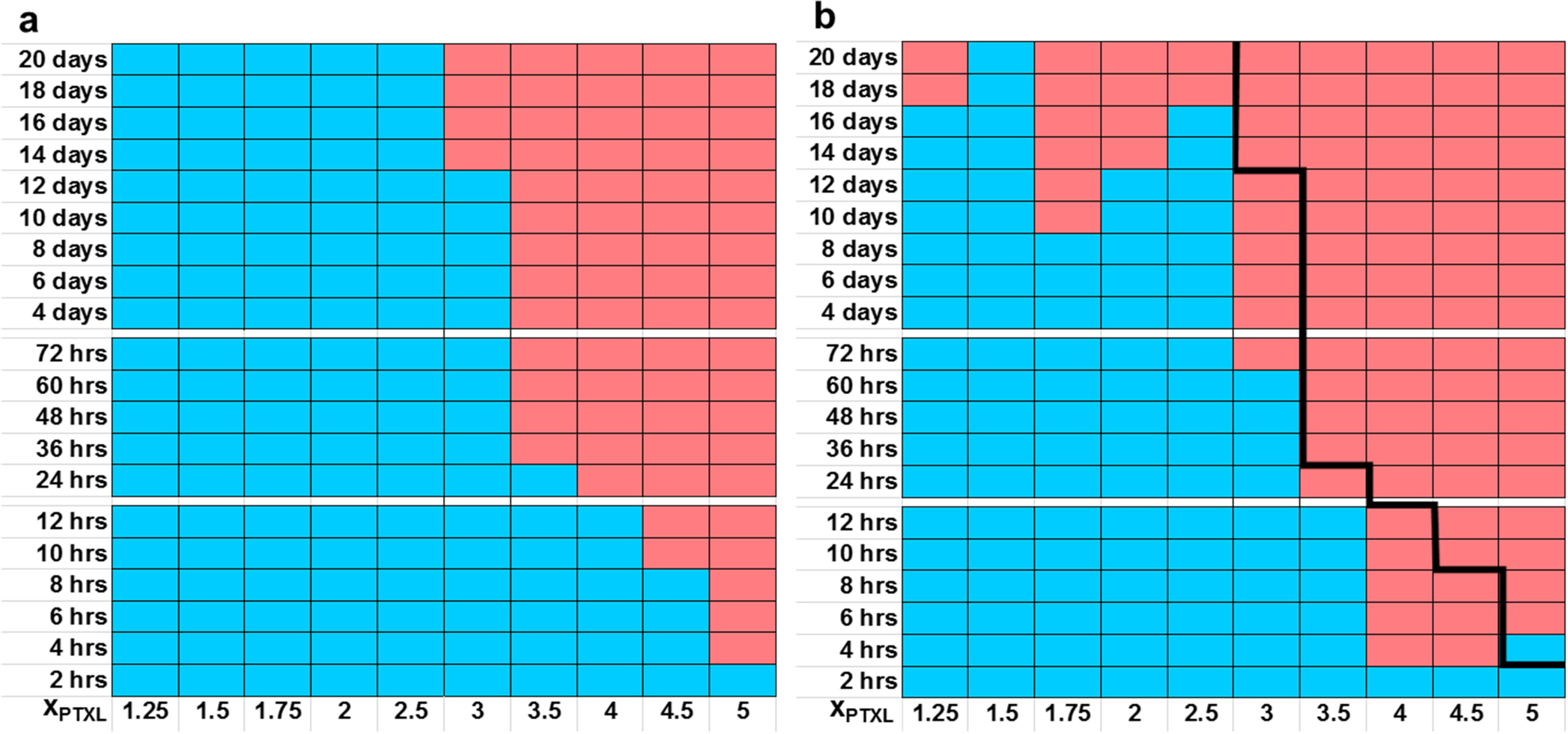
Kinetic phase diagrams of PTXL solubility in CL_PTXL_ NPs prepared from DOTAP, DOPC and PTXL (30:70–x_PTXL_:x_PTXL_ mole ratio). DIC microscopy (see Figure 2) was used to assess whether PTXL crystallization had occurred at the indicated times after hydration. Blue color indicates absence of PTXL crystals (PTXL remained soluble in the membranes), while pink color indicates presence of PTXL crystals. (a) Stability of PTXL in *unsonicated* liposomes. (b) Stability of PTXL in *sonicated* liposomes. The black line, showing the solubility boundary for unsonicated liposomes, is included as a reference to facilitate comparison.

The phase diagrams show that PTXL is relatively unstable within membranes when it is incorporated at > 3 mol% in DOTAP/DOPC liposomes, phase separating on a time scale of hours to one day. In contrast, unsonicated samples incorporating PTXL at < 3 mol% PTXL did not phase separate within the timeline of observation (with selected samples monitored over 30 days). At 3 mol% PTXL content, PTXL-loaded CLs display moderate stability, which is reduced if the sample was subjected to sonication. Indeed, as evident from a comparison of Figure 3a (unsonicated) to Figure 3b (sonicated), sonication reduced the stability of CL_PTXL_ particles over the entire range of PTXL contents investigated. (The black line in Figure 3b marks the boundary denoting onset of crystal formation in unsonicated samples.)

The sonicated, smaller CL_PTXL_ NPs have a much higher curvature (C ≈ 1/200 nm^−1^ versus ≈ 1/800 nm^−1^ for unsonicated PTXL-loaded CLs), which appears to promote faster nucleation and growth rates. A possible rationale for this observation is that high membrane curvature breaks the symmetry between the inner and outer lipid monolayers (i.e., C_monolayer_^outer^ > 0 while C_monolayer_^inner^ < 0 for small NPs, whereas larger, nearly flat unsonicated membranes (C ≈ 0) have very small differences in curvature between the monolayers). This would lead to preferred PTXL partitioning into one monolayer in small NPs and thus a higher local PTXL concentration and faster nucleation time.

#### Synchrotron Small Angle X-Ray Scattering (SAXS)

We used SAXS to quantitatively confirm the existence of phase-separated PTXL crystals (observed in DIC) by their signature diffraction peaks [50–52]. We studied selected CL_PTXL_ NPs and DOTAP/DOPC/PTXL multilayers condensed with DNA by SAXS at different times after liposome hydration for comparison with the kinetic phase diagram results obtained by DIC and to gather detailed structural information about PTXL-containing membranes.

The top profile in Figure 4a depicts SAXS from a control sample of CLs (DOTAP:glycerol monooleate (GMO), 50:50 mole ratio) without PTXL, where only weak form factor scattering from the lipid membranes is observed. The bottom profile shows SAXS from CL_PTXL_ NPs (DOTAP:GMO:PTXL, 50:44:6 mole ratio) where three strong diffraction peaks arise from the presence of phase-separated PTXL crystals. These crystals are also optically visible in the x-ray capillary. The three characteristic PTXL diffraction peaks are located at *q*_P1_ = 0.291 Å^−1^, *q*_P2_ = 0.373 Å^−1^, and *q*_P3_ =0.436 Å^−1^ in the low-*q* range (0.005 Å-1 < *q* < 0.5 Å^−1^) probed in our SAXS experiment.

**Figure 4.**
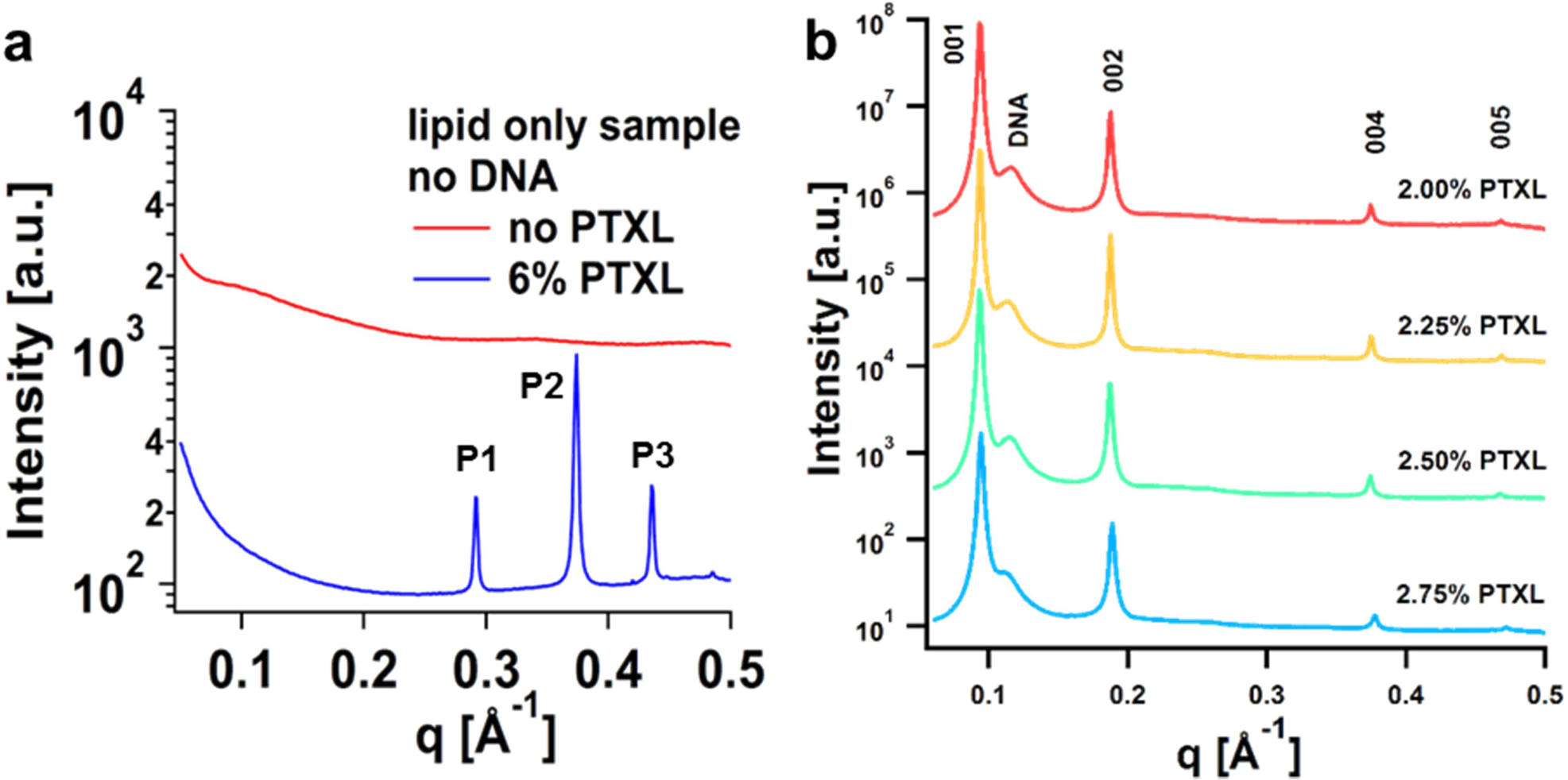
X-ray scattering analysis of CL_PTXL_ NPs. (a) Representative examples of the average scattering intensity for uncondensed (DNA-free) lipid samples without PTXL (red) and CL_PTXL_ NPs with high PTXL content (blue). Only the scattering form factor appears for the lipid sample when the lipid bilayers are not condensed. When PTXL has crystallized (blue curve), peaks due to PTXL crystals are observed at P1 (*q*=0.291 Å^−1^), P2 (*q*=0.373 Å^−1^), and P3 (*q*=0.436 Å^−1^). (b) Small angle x-ray scattering intensity for four CL_PTXL_ NP samples with high PTXL solubility (PTXL content < 3 mol%; DOTAP/DOPC/PTXL, 30:70–x_PTXL_:x_PTXL_ mole ratio) that were condensed with DNA on the fourth day after hydration. In these samples, PTXL remains soluble (no P1, P2 or P3 peaks are observed). However, because the samples have been condensed with DNA, the (001), (002), (004), and (005) peaks characteristic of the L_α_^C^ phase appear, along with the broad DNA–DNA correlation peak, as labeled.

To enable detailed structural analysis of PTXL-loaded membranes, we had to overcome the problem of weak scattering from the dilute CL samples (Figure 4a). Thus, we concentrated CL_PTXL_ NPs by complexing them with oppositely charged macromolecules, using anionic DNA as the condensing agent. The resulting CL_PTXL_–DNA complexes can be further compacted into a high-membrane-concentration pellet by centrifugation. We verified that CL_PTXL_–DNA complexes had equivalent human cancer cell death efficacy compared to CL_PTXL_ NPs. In the absence of PTXL, SAXS has shown that CL–DNA complexes form the lamellar L_α_^C^ phase for CLs composed of DOTAP and DOPC [53–56]. The multilayered structure of the L_α_^C^ phase, consisting of onion-like NPs with diameter ≈300–400 nm, is also directly observed in cryoTEM [57,58]. Figure 4b shows SAXS profiles for CLPTXL–DNA complexes (DOTAP:DOPC:PTXL 30:70–x_PTXL_:x_PTXL_ mole ratio; lipid/DNA charge ratio = 1.5) at x_PTXL_ = 2.75, 2.50, 2.25, and 2.00, prepared on the fourth day after liposome hydration. The observed sharp scattering peaks correspond to the (00L) peaks (L = 1,2,4, and 5) of the L_α_^C^ phase with interlayer spacing d_lamellar_ = 2π/*q*_001_ = 67 Å consisting of the combination of the thickness of the lipid bilayer (containing DOTAP, DOPC, and PTXL) and the water layer (containing a layer of DNA) [53–56]. The (003) peak is not observed because it is close to a minimum of the x-ray form factor of the CL_PTXL_–DNA complexes. The broader shoulder peak to the right of the (00 1) peak at *q*_DNA_ = 0.112–0.116 Å^−1^ is due to DNA–DNA correlations and yields an average DNA interaxial spacing *d*_DNA_ = 2π/*q*_DNA_ = 54.0–55.9 Å [53]. The scattering from these samples containing < 3 mol% PTXL (Figure 4b) shows no evidence of PTXL-related peaks, consistent with the kinetic phase diagram (Figure 3b).

Figure 5 shows data from time-dependent SAXS studies. For each CL_PTXL_ NP sample, aliquots were freshly complexed with DNA at each time point (to avoid potential artifacts resulting from the confinement of the membranes in the CL_PTXL_–DNA complexes) and assessed by SAXS to check for the signature PTXL crystal diffraction peaks. For samples incorporating larger amounts of PTXL (x_PTXL_ = 3, 3.5, and 4.5), the PTXL diffraction peaks appeared on or before day 4 of the experiment. This is illustrated by Figure 5 (a and b), which shows the scattering profiles at four consecutive days after liposome hydration (labeled D1 through D4). The appearance of the P1 and P3 peaks is the primary indication of PTXL crystal formation for this set of data because the positions of the P2 and (004) peaks are very close together. The PTXL peaks appear on day 4 for 3.0 mol% PTXL, day 3 for 3.5 mol%, and day 2 for 4.5 mol% PTXL. All peak positions for the samples in Figure 4b and Figure 5 are reported in the Supporting Information (Table S1).

**Figure 5.**
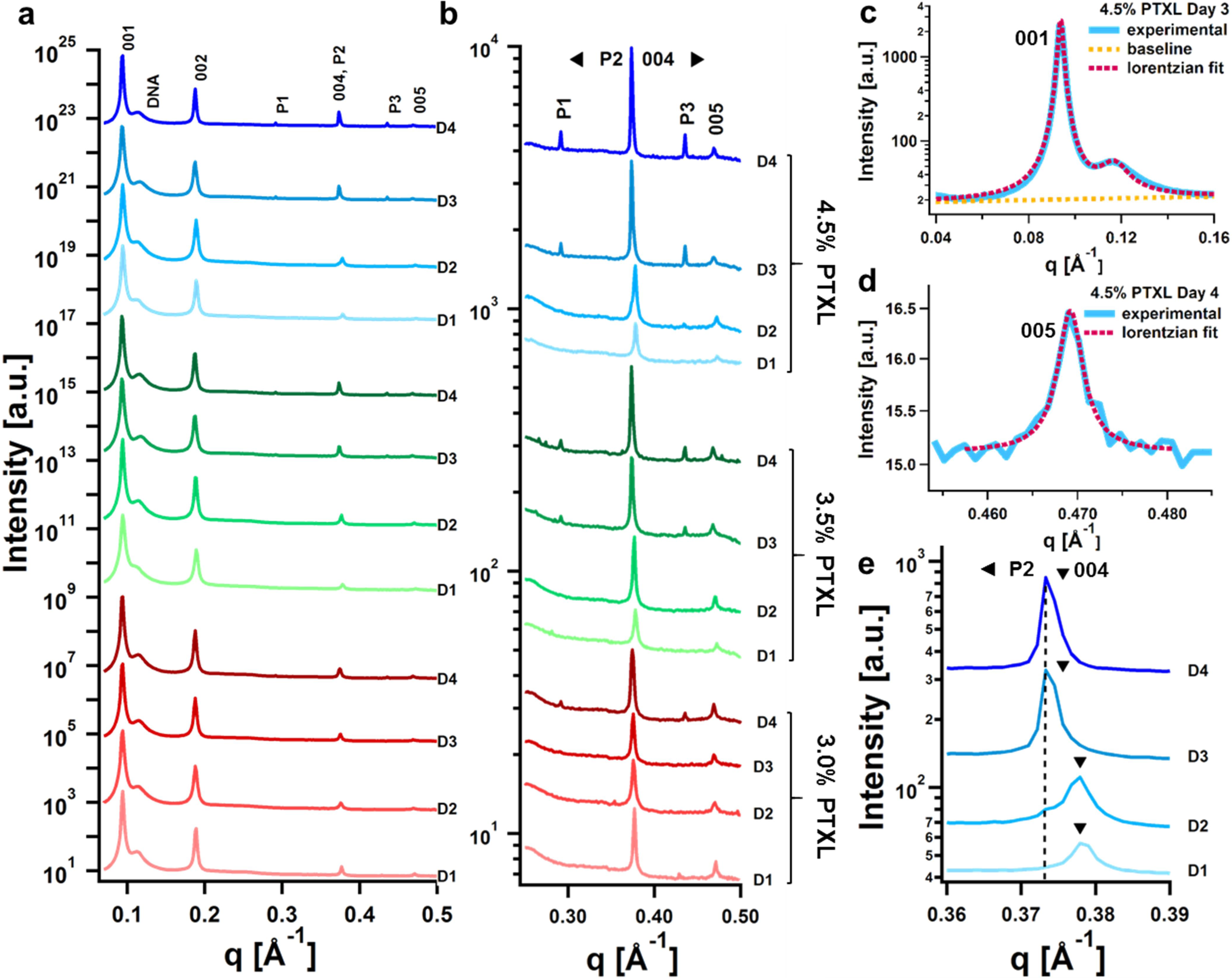
X-ray scattering analysis of PTXL crystallization from CL_PTXL_ NPs. Scattering intensity is plotted for samples containing 3.0 (red lines), 3.5 (green lines), and 4.5 (blue lines) mol% PTXL (DOTAP/DOPC/PTXL, 30:70–x_PTXL_:x_PTXL_ mole ratio). Over the course of 4 d following hydration, aliquots of the CL_PTXL_ NPs were condensed with DNA each day (day of scan designated on graphs as D1–D4) and formation of PTXL crystals was assessed with SAXS. (a) Full scattering spectra for all samples over the investigated range of the scattering vector *q* (*q*=0.05–0.5 Å^−1^). Peaks originating from the stacked membranes of the lamellar L_α_^C^ structure are labeled as 001, 002, 004, and 005, while the DNA–DNA correlation peak is marked DNA, and P1, P2, P3 mark peaks originating from PTXL crystals (see Figure 4). (b) Expanded view of the plots in (a) around the PTXL peaks (*q*=0.25–0.50 Å^−1^). The appearances of peaks at P1 and P3 are the clearest indication of the presence of PTXL crystals. (c) An example of peak fitting for the (001) and *q*_DNA_ peaks. The dashed line indicates a background-subtracted fit as the sum of two Lorentzian functions. The background scattering is shown by the orange dashed line with the form *I^bg^*(*q*) = *m*\**q* + *y*_0_. Each Lorentzian function was written as *S*(*q*) = *A*/[κ^2^ + (*q*-*q_0_*)^2^)], where *q*_0_ and κ correspond to the peak position and the hwhm, respectively. For the (001) SAXS peak, *A*_001_ = 3.80 × 10^−3^, *q*_001_ = 0.09374 Å^−1^, κ_002_ = 1.16 × 10^−3^ Å^−1^; for the *q*_DNA_ SAXS peak, *A*_DNA_ = 1.36 × 10^−3^, *q*_DNA_ = 0.1168 Å^−1^, κ_DNA_ = 6.64 × 10^−3^ Å^−1^. (d) Example fit for the (005) peak, which was used to determine the interlayer lamellar distance (dictated by the membrane bilayer thickness). The (005) peak was fit to a single Lorentzian with a constant background scattering of *y_0_* = 15.1, where *A*_005_ = 3.97 × 10^−6^, *q*_005_ = 0.4692 Å^−1^, and κ_005_ = 1.70 × 10^−3^ Å^−1^. In this particular series of samples (30 mol% DOTAP, DOTAP/DNA charge ratio of 1.5), the prominent P2 peak overlaps the (004) peak. (e) Expanded view of the region around the P2 and (004) peaks (4.5 mol% PTXL, days 1-4). The black arrows indicate the predicted position of the (004) peak for each sample, based on the position of the (005) peak. As PTXL leaves the membrane, the (004) peak shifts to lower q (corresponding to a thickening of the membrane), toward the P2 peak (position indicated by dashed line) which appears as the PTXL crystals form.

Figure 5c shows a representative lineshape analysis for the (001) and *q*_DNA_ peaks (x_PTXL_ = 4.5, day 3). To get an accurate value for the peak positions, these overlapping peaks were fit simultaneously as the sum of two Lorentzian functions (dashed line). Each Lorentzian function was written as *S*(*q*) = *A*/[κ^2^ + (*q*−*q*_0_)^2^],with *q*_0_, κ and *A* as the fit parameters. The parameters *q*_0_ and κ correspond to the peak position and the half-width at half-maximum, respectively, while *A*/κ^2^ is the peak intensity. The background scattering (orange line) was approximated by a sloped line with the form *I^bg^*(*q*) = *m*\**q* + *y*_0_.

Figure 5d shows a representative example fit for the (005) peak (for x_PTXL_ = 4.5, day 4), which was used to determine the lamellar interlayer spacing *d*_lamellar_ = 2π/(*q*_005_/5). The (005) peak was used to measure *d_lamellar_* because it is the highest order diffraction peak in the q-range of our SAXS study. It is expected to show the largest variation in peak position, thus yielding the most accurate measurement of changes in *d_lamellar_*. The (005) peak was fit to a single Lorentzian with a constant background. As evident in Figures 5c and 5d, the Lorentzian fits show good agreement with the experimental scattering data, enabling an accurate measurement of peak positions.

Figure 5e shows enlarged sections of the scattering data for the sample containing 4.5 mol% PTXL in the vicinity of the (004) peak on days 1 through 4. The arrowheads in Figure 5e point to the position of q004 as deduced from the position of *q*_005_ (which has no nearby peak). On days 1 and 2, when PTXL is still soluble in the membrane, *q*_004_ = 0.378 Å^−1^, but on days 3 and 4, when PTXL has largely phase-separated into crystals (i.e., when the P1 and P3 diffraction peaks are strong), *q*_004_ = 0.375 Å^−1^. This indicates a change in membrane thickness upon PTXL crystallization (see below). The onset of P2 at a slightly lower *q* (0.373 Å^−1^, indicated by the dashed line) than *q*_004_ gives rise to a closely spaced doublet peak, which is quite evident from the asymmetric shape of the peak at day 3. The P2 and P3 peaks emerge on day 2 (see the small bump at the dashed line marking P2 in Figure 5e and the small P3 peak in Figure 5b (4.5% PTXL) on D2), but based on their small size and the unchanged position of *q*_004_ (from day 1), most of the PTXL remains soluble in the membrane.

Figure 6a displays plots of the interlayer spacing, *d*_lamellar,_ calculated from the 5^th^ harmonic (*d*_lamellar_ = 2π/(*q*_005_/5)), as a function of time (day 1 through day 4) for the samples with x_PTXL_ = 3, 3.5, and 4.5 mol% (DOTAP/DOPC/PTXL, 30:70–x_PTXL_:x_PTXL_ mole ratio). The data shows an increase in *d*_lamellar_ from days 1 and 2 (when all PTXL is still soluble in the membranes) to days 3 and 4 (when insoluble PTXL crystals are present). An increase in *d*_lamellar_ implies an increase in the average membrane thickness because the thickness of the water layer containing the monolayer of DNA, electrostatically adhered to neighboring cationic membranes, is nearly constant at around 2.5 nm [53]. This increase in membrane thickness on days 3 and 4 (with PTXL crystals present) is consistent with depletion of PTXL from the membranes (i.e., replacing a DOPC molecule with a shorter PTXL molecule (see Figure 1b) is expected to thin the membrane; conversely, membranes thicken as the molar ratio of PTXL to DOPC and DOTAP decreases).

**Figure 6.**
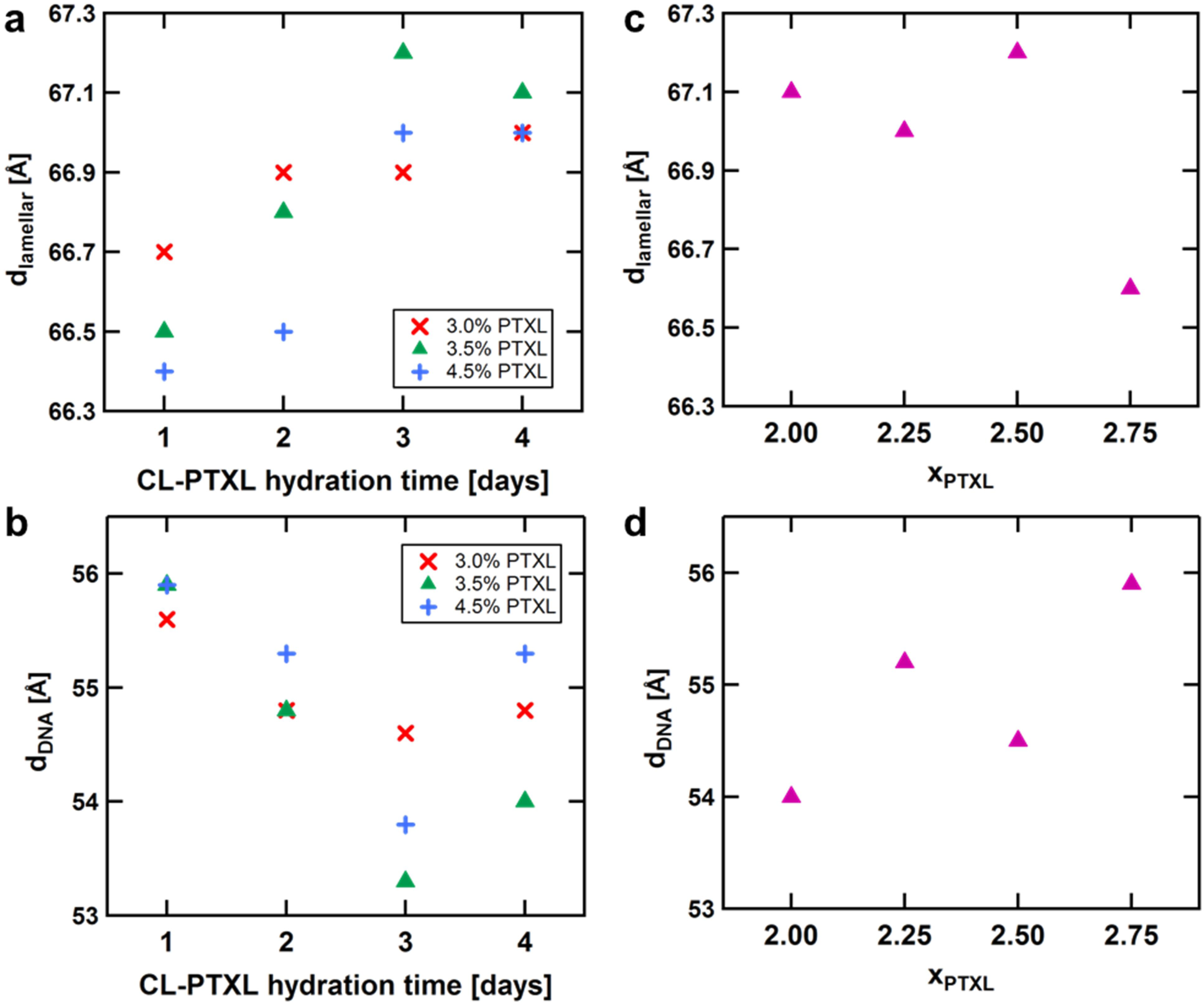
Lamellar interlayer distance and DNA interaxial spacing for DOTAP/DOPC/PTXL samples (30:70–x_PTXL_:x_PTXL_ mole ratio) condensed with DNA as derived from x-ray lineshape analysis. (a,b) Plots of *d*_lamellar_ (a) and *d*_DNA_ (b) as a function of time for samples which exhibited PTXL crystallization. (c,d) Plots of *d*_lamellar_ (c) and *d*_DNA_ (d) against PTXL content for the stable samples (no PTXL crystallization; < 3 mol% PTXL) on day 4. The lamellar interlayer distance *d*_lamellar_ is the sum of the thicknesses of the lipid bilayer and the DNA-containing water layer. It was calculated from the position of the (005) peak as *d*_lamellar_ = 2π/(*q*_005_/5). The DNA–DNA spacing was calculated from the DNA–DNA correlation peak (*d*_DNA_=2π/*q*_DNA_). See text for discussion.

We also observed that the DNA interaxial spacing (*d*_DNA_) decreases with time as PTXL crystals form on days 3 and 4 and PTXL is depleted from the membrane (Figure 6b). This decrease in *d*_DNA_ arises because the charge density of DOTAP/DOPC/PTXL membranes increases upon the loss of neutral PTXL from the membrane (i.e., the number of charged molecules in the membrane (DOTAP) remains constant, but the area of the membrane is reduced by the loss of PTXL). An increase in cationic membrane charge density is known to drive a decrease in *d*_DNA_ because the decrease in *d*_DNA_ increases the anionic charge density (as required to maintain overall local charge neutrality between cationic membranes and anionic DNA) [53,54,56].

Figure 6c depicts the interlayer spacing as a function of increasing PTXL content for the samples where PTXL did not phase separate (see Figure 4b), i.e., for xPTXL = 2, 2.25, 2.5, and 2.75 (DOTAP/DOPC/ PTXL, 30:70–x_PTXL_:x_PTXL_ mole ratio). The data shows the expected decrease in *d*_lamellar_ (and thus the membrane thickness) as increasing amounts of DOPC are replaced by the shorter PTXL molecule (consistent with the behavior found in Figure 6a). Figure 6d shows that the DNA interaxial spacing increases when the PTXL content in the membrane is increased, similar to the behavior seen in Figure 6b. In this case, the number of molecules in the membrane is constant (DOPC is replaced with PTXL to increase the PTXL content). However, replacing a DOPC molecule with a shorter but thicker PTXL molecule (see Figure 1b) leads to an increase in the average distance between lipids (increase in the lateral area per lipid) and a lower membrane charge density. This lowering of the membrane charge density with increasing PTXL content drives the increase in DNA spacing (i.e., lowering of the anionic charge density) as noted above.

This data shows that our new methodology, of using cationic membranes complexed with oppositely charged DNA to produce highly condensed aggregates suitable for *in situ* high-resolution synchrotron SAXS studies, has enabled us to accurately detect and measure small variations in the membrane interlayer spacing and the DNA interaxial spacing due to changes in the relatively small amount of PTXL (< 5 mol%) incorporated in the cationic membranes as a function of time. After probing the time scale of PTXL solubility in membranes using optical microscopy and SAXS, we set out to correlate the observed differences between formulations incorporating varied amounts of PTXL to their drug delivery efficacy as measured by their toxicity to human cancer cell lines in vitro.

#### Cell Viability Characterization of CL_PTXL_ NP Efficacy

To begin our investigations of the efficacy (i.e. ability to induce cancer cell death) of CL-based PTXL carriers, we measured the approximate IC-50 (the drug concentration achieving half the maximal effect) for cytotoxicity in two human cancer cell lines. (We found that the exact IC-50 varies with cell passage number.) We obtained this baseline value using CLs prepared from DOTAP:DOPC:PTXL at a molar ratio of 50:47:3 (to mimic the proprietary EndoTAG-1 formulation) [59]. The plots of cell survival (normalized to untreated cells) as a function of increasing PTXL concentration are shown in Figure S1 in the Supporting Information. For PC3 cells, (prostate cancer metastasis) the IC-50 ≈ 20 nM and the cell survival curve exhibits a steep slope (between 10 and 50 nM). For the M-21 cell line (melanoma metastasis), IC-50 ≈ 45 nM with a more gradual slope (spanning the range of 5–200 nM).

Next, we compared the efficacy of CL-based PTXL carriers of varied composition side-by-side at the predetermined IC-50 (thereby eliminating errors that can occur when using cells of different passage number). For these experiments, we chose CL_PTXL_ NPs at PTXL contents covering the three different regimes of PTXL membrane solubility observed in the kinetic phase diagram (Figure 3): long-term solubility (< 3 mol% PTXL), moderate solubility (3 mol% PTXL), and low solubility (> 3 mol% PTXL). To allow a valid comparison of cell survival for NPs with different PTXL content, the total applied concentration of PTXL was fixed near the IC-50 value. Thus, CL_PTXL_ NPs with higher PTXL content in the membrane yielded correspondingly lower final molar concentrations of lipid.

Figure 7 depicts PC3 and M21 cell survival in response to 20 nM and 60 nM PTXL concentration, respectively, as a function of increasing PTXL content for CL_PTXL_ NPs with membrane composition DOTAP:DOPC:PTXL=50:50-x_PTXL_:x_PTXL_ (molar ratio). For both cell lines, the data reveals a surprising nonmonotonic dependence on the PTXL content. The most effective compositions, with the lowest cell survival, are the formulations with PTXL content below 3 mol%. Cell survival is higher for 3 to 7 mol% PTXL, and then decreases again above 7 mol%. The more sensitive response of PC3 cells to PTXL concentration (steeper slope of the cell survival curve, see Figure S1) may explain the more dramatic differences in cell survival for PC3 cells compared to the more incremental differences observed for M21 cells.

**Figure 7.**
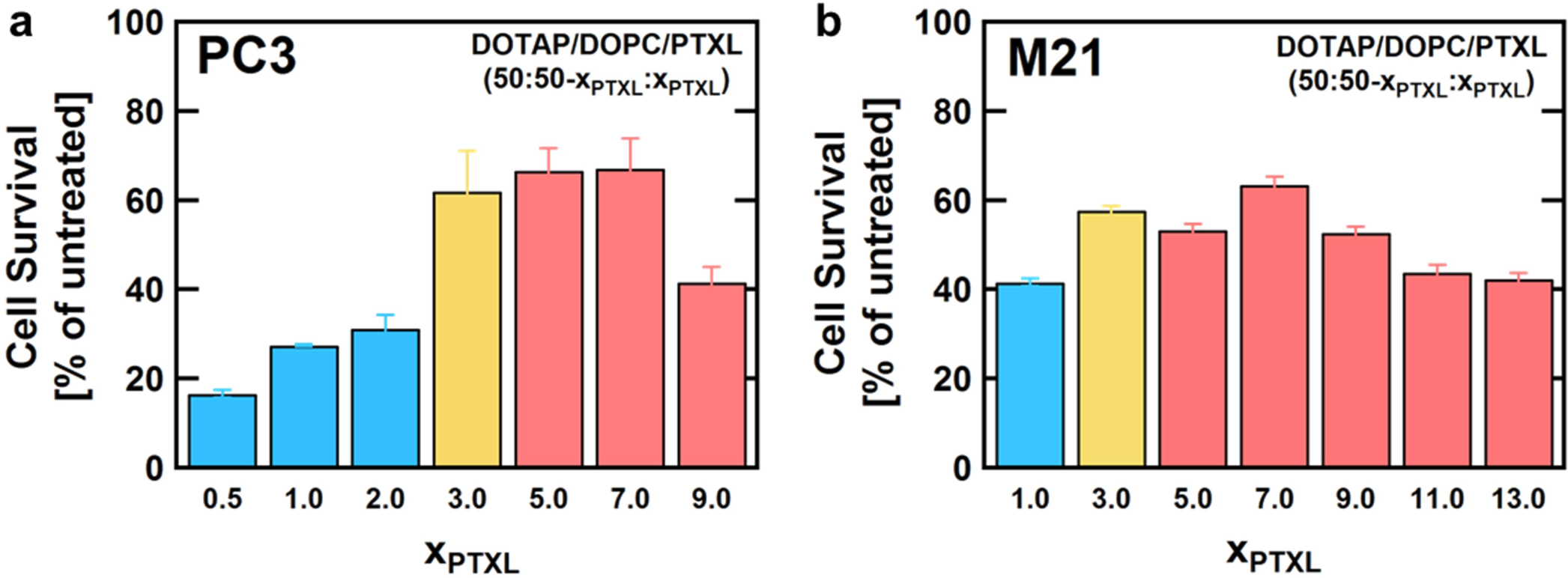
Cell survival data for CL_PTXL_ NPs of varied PTXL content in two human cancer cell lines. The total concentration of PTXL applied to cells was kept constant at 20 nM for PC3 cells (a) and 60 nM for M21 cells (b). These concentrations are near the IC-50s for PTXL in EndoTAG-1-like NPs, which were determined in previous experiments to assess the sensitivity of each cell line to PTXL. Because the total amount of applied PTXL was constant, the amount of applied lipid decreased as the PTXL content increased. Cell viability was measured 72 h after the NPs were added to cells and is normalized to that of untreated cells.

To control for lipid toxicity—especially for CL_PTXL_ NPs at low PTXL content, where the most lipid was administered to the cells—we measured the viability of PC3 and M21 cells incubated with increasing concentrations of DOTAP/DOPC (50:50, mol:mol) small unilamellar CLs. As evident from the data shown in Figure 8, no toxicity is observed even at 100 μM total lipid, the upper limit tested in this control experiment. This is consistent with literature values for various lipids, which showed toxicity arising in the 100-200 μM range for the most toxic lipids in the study [60], while toxicity does not set in until 3000–4000 μM for the common neutral lipids mixed-chain phosphatidylcholine (egg lecithin) and dipalmitoylphosphatidylcholine (DPPC). The final total lipid concentration in our experiments probing CL_PTXL_ NP toxicity never exceeded 6 μM lipid, and therefore the observed toxicity is due to the delivered PTXL.

**Figure 8.**
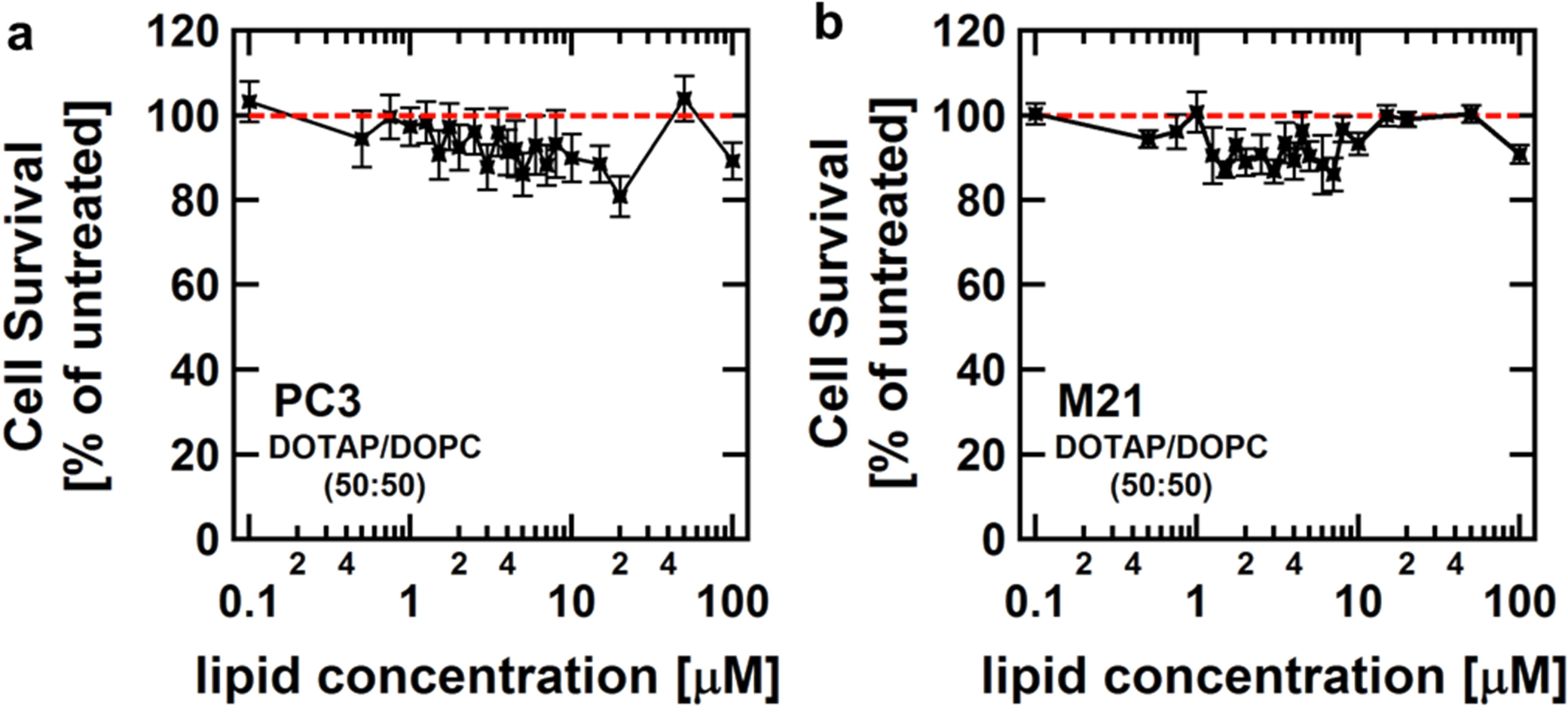
Assessment of lipid toxicity in a) PC3 cells (human prostate cancer) and b) M21 cells (human melanoma). Liposomes consisting of DOTAP and DOPC (1:1 mole ratio) were diluted in DMEM from 100 to 0.1 μM total lipid and added to cells. Cell viability was measured 72 h after the liposomes were added to the cells and is normalized to that of untreated cells. No lipid toxicity is apparent in this concentration range.

To verify the unexpected nonmonotonic trend of toxicity as a function of PTXL content, we expanded on the limited snapshot of information afforded by the experiments reported in Figure 7. Thus, we measured the IC-50 for cell toxicity of CL_PTXL_ NPs at PTXL contents of 1, 3, and 9 mol% for PC3 and M21 cells (Figure 9). In these experiments NPs were added to cells within 2–3 hours after hydration. For both cell lines, efficacy (diminishing of cell survival) is greatest when PTXL is incorporated at only 1 mol% (and thus more lipid is used as a carrier). The IC-50 for CL_PTXL_ NPs with 1 mol% PTXL is more than a factor of 2 lower than that for carriers with 3 mol% PTXL, both for PC3 (Figure 9a) and M21 (Figure 9b) cells. The cell survival curves for CL_PTXL_ NPs at both 1 and 9 mol% PTXL are left-shifted with respect to the baseline formulation of 3 mol% PTXL, consistent with the bell-shaped curves shown in Figure 7. Thus, this data further supports the presence of two optimal activity regimes at low (≈ 1 mol%) and high (≈ 9 mol%) PTXL content.

**Figure 9.**
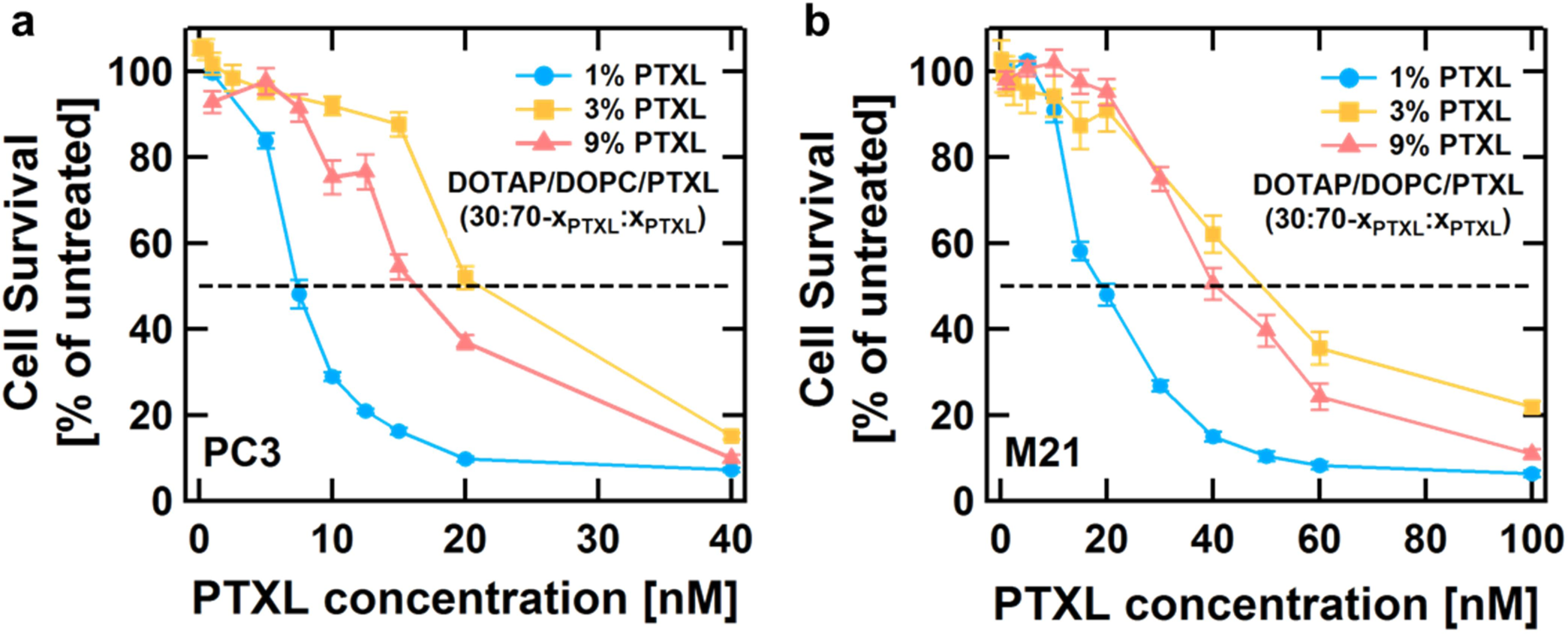
Cell survival as a function of PTXL concentration for CL_PTXL_ NPs containing 1, 3 or 9 mol% PTXL added to a) PC3 cells and b) M21 cells. The dashed line indicates 50% cell death. For PC3 cells, the IC-50 is highest (about 20 nM) for 3 mol% PTXL, and lower for both 1 mol% PTXL (about 7.5 nM) and 9 mol% PTXL (about 15 nM). Similarly, the IC-50 in M21 cells is highest (about 50 nM) for 3 mol% PTXL, but lower for 1 mol% PTXL (about 20 nM) and 9 mol% PTXL (about 40 nM).

Our kinetic phase diagram study makes it evident that the microscopic states of PTXL in membranes evolve with time spent in the aqueous milieu (i.e., from a mixed state to a demixed state of PTXL nucleation and growth). Thus, we also investigated the efficacy of CL_PTXL_ NPs with PTXL content of 1 and 9 mol% as a function of time after hydration. The data from these experiments (Figure 10) demonstrates that cells respond very differently to the same dose of PTXL depending on whether or not the PTXL is still soluble within the CL membranes. When more lipid is used to solubilize PTXL (1 mol% PTXL content), efficacy remains high (low cell survival) irrespective of time after hydration. In contrast, a significant drop in efficacy occurs (cell survival increases) over time for the CL_PTXL_ NPs at 9 mol% PTXL content (which are more efficient than NPs containing 3 mol% PTXL at earlier times: see Figures 7 and 9). NPs with 3 mol% PTXL show a similar trend to those with 9 mol% PTXL, losing efficacy over time.

**Figure 10.**
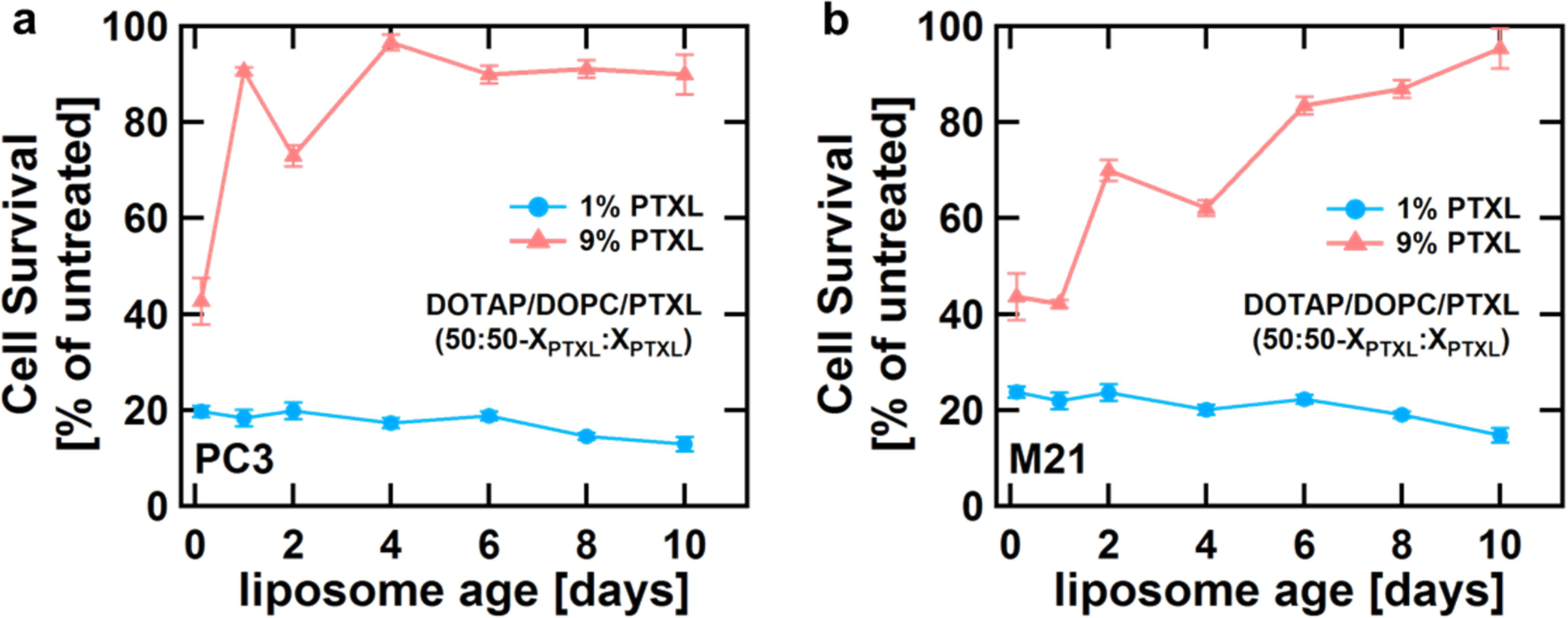
Cell survival data for CL_PTXL_ NPs with selected PTXL content in two human cancer cell lines as a function of time after hydration in water. The total concentration of PTXL applied to cells was kept constant at 22.5 nM for PC3 cells (a) and 65.0 nM for M21 cells (b). The CL_PTXL_ NPs containing 1 (blue) or 9 (pink) mol% PTXL (50 mol% DOTAP) were hydrated and sonicated at different time points to be applied to the cells at the same time. Cell viability was measured 72 h after the NPs were added to cells and is normalized to that of untreated cells.

The improved activity of carriers with well-solubilized PTXL (1 mol% PTXL) can be readily rationalized by the fact that the drug remains bioavailable for longer. The toxicity of PTXL requires about 72 hours to take full effect, and the properties of these carriers ensure that the drug will not phase separate in this period. Conversely, the drug delivery efficacy that reemerges at higher PTXL loading, at 9 mol% PTXL, is highly unexpected. It may be that at this concentration, with PTXL supersaturated in the membrane, PTXL molecules are readily expelled from the lipid membranes soon after hydration. This PTXL may diffuse directly into the cell membranes, which appear as a “lipid sink.” As time progresses and the PTXL phase separates and crystallizes (becoming biologically inert), it would render these carriers less and less effective.

## Conclusions

DIC optical microscopy and high-resolution synchrotron SAXS have allowed us to map the kinetic phase behavior of DOTAP/DOPC-based CL carriers of PTXL and correlate distinct stability and efficacy regimes. To date, all efficacy studies of liposome-based PTXL carriers (including in animal models and cancer chemotherapy clinical trials) have incorporated PTXL in lipid NPs near the membrane solubility limit at 3 mol% [33,34,39,59]. In strong contrast, the work reported here shows that for NP delivery shortly after preparation (on the time scale of hours) two different PTXL composition regimes, below and above 3 mol% PTXL, have notably higher efficacy in prostate (PC3) and melanoma (M21) human cancer cells. A further significant finding of our study is that the efficacy of CL_PTXL_ NPs with higher PTXL content (≥ 3 mol%) is strongly dependent on the time between NP preparation and delivery. The unique efficacy (on short time scales) of CLs with PTXL content of 9 mol% is worth future consideration. These liposomes require much less lipid ‘solvent’ for delivery—a benefit should lipid off-target toxicity or cost become a concern. While not well-suited for long-term circulation, this carrier may be a better candidate when the use of local injection is appropriate to achieve a high initial drug concentration.

Our experiments suggest that DOTAP/DOPC liposomes loaded with 3 mol% PTXL are still likely to phase separate on a time scale of days. This was observed in plain water, and would likely be accelerated in a biological environment. To our knowledge, it has not yet been explored if liposomes encapsulating PTXL below its solubility limit can improve pharmacokinetics, pharmacodynamics, or biodistribution *in vivo*. It may be counterintuitive to use particles below their nominal drug-loading capacity, but our data suggests that this may facilitate actual drug targeting and enhance therapeutic outcomes for hydrophobic drugs by improving drug retention, as long as the threshold of lipid toxicity is not exceeded.

Finally, we presented a new methodology involving in situ synchrotron SAXS of PTXL-loaded cationic multilamellar membranes condensed by DNA, which has allowed us to directly confirm the presence of PTXL in membranes and to observe the time-dependent depletion of PTXL from membranes upon PTXL crystal formation by measuring small variations in the membrane interlayer and DNA interaxial spacings. We expect this nondestructive approach to be generally applicable to other hydrophobic molecules. Besides allowing for precise measurements of the dimension of PTXL-containing membranes under realistic, bulk-water conditions, complexes of such membranes with DNA (or siRNA) constitute a promising class of dual-cargo delivery vehicles which may be used to deliver nucleic acids to work synergistically with the delivered PTXL [31].

## Acknowledgment

This work was supported by the National Institute of Health under award GM-59288. The work was also supported in part by the National Science Foundation under award DMR-1401784 (phase behavior of cationic membranes containing PTXL). VS was supported by the National Science Foundation Graduate Research Fellowship Program under Grant No. DGE 1144085.

## Supporting Information

PDF includes experimental data for an IC-50 determination for each cell line (M21 and PC3), as well as all peak positions for x-ray scattering data shown in Figures 4b and 5.

